# Molecular strategies for antibody binding and escape of SARS-CoV-2 and its mutations

**DOI:** 10.1101/2021.03.04.433970

**Authors:** Mohamed Hendy, Samuel Kaufman, Mauricio Ponga

## Abstract

The COVID19 pandemic, caused by SARS-CoV-2, has infected more than 100 million people worldwide. Due to the rapid spreading of SARS-CoV-2 and its impact, it is paramount to find effective treatments against it. Human neutralizing antibodies are an effective method to fight viral infection. However, the recent discovery of new strains that substantially change the S-protein sequence has raised concern about vaccines and antibodies’ effectiveness. Here, we investigated the binding mechanisms between the S-protein and several antibodies. Multiple mutations were included to understand the strategies for antibody escape in new variants. We found that the combination of mutations K417N and E484K produced higher binding energy to ACE2 than the wild type, suggesting higher efficiency to enter host cells. The mutations’ effect depends on the antibody class. While Class I enhances the binding avidity in the presence of N501Y mutation, class II antibodies showed a sharp decline in the binding affinity. Our simulations suggest that Class I antibodies will remain effective against the new strains. In contrast, Class II antibodies will have less affinity to the S-protein, potentially affecting these antibodies’ efficiency.

## 1 Introduction

The severe acute respiratory syndrome coronavirus 2 (SARS-CoV-2) enters host cells via a chemo-mechanical attachment to the human cells. The attachment is aided by the human angiotensin-converting enzyme 2 (ACE2) receptor, which is usually expressed in epithelial cells found in blood vessels, lungs, and the nasal cavity (*1*). After infecting host cells, SARS-CoV-2 could develop into severe pneumonia with the potential to cause severe injuries and death (*2*). With more than 100 million infections and 2.2 million deaths worldwide, and cases still rising, it is imperative to find treatments that can provide long-lasting immunity to COVID19.

SARS-CoV-2 uses a transmembrane spike (S) glycoprotein forming homotrimers on the virus’ capsid (*1*, *3*) to attach to the receptors. The S-protein has two functional subunits, nominally named S1 and S2. A key characteristic feature of SARS-CoV-2 is that in its prefusion stage, the S1 subunit has two states labeled as *up* and *down*, whereby the S1 subunit is exposed and retracted, respectively (*4*). The *up* level serves as the basis to expose the receptor-binding domain (RBD) in the S1 protein for binding to the receptor. By way of contrast, the *down* state of the S1 protein serves as a shield for the spike against antibodies.

The S1 subunit, and, more specifically, the RBD motif (residues 417-505 of the virus’ sequence), is directly responsible for the attachment to the cell receptors. The RBD strongly links to the ACE2 receptors with a dissociation constant measured between *k_D_* = 1.2 − 22 nM (*1, 3, 5, 6*). After the initial attachment, the S-protein is cleaved at two locations. This process is thought to be assisted by furin (*7*). Subsequently, the S-protein undergoes large conformational changes to expose its fusion peptide to diffuse through the cell’s lipid membrane for posterior fusion (*8*). It is known that SARS-CoV-2 enters cells via at least two different mechanisms (*9, 10*), i.e., via endocytosis (*9, 11*–*13*), or direct fusion to the cell membrane assisted by the transmembrane protease serine 2 (TMPRSS2) (*9*). However, several key steps in this process remain unknown (*14*). Due to its importance to enter cells, several vaccines target the S-protein as a primary candidate.

In response to this foreign entity, the human’s immune system develops antibodies that link to several parts of the virus to stop and deter the viral infection. Recent studies on sera of convalescent patients have shown a large variety of effective human neutralizing antibodies (hNAbs) for SARS-CoV-2 targeting the S-protein, particularly its RBD (*15, 16*). For instance, Barnes and co-workers (*17*) have classified hNABs relative to their binding strategy and have elucidated four antibody classes. Class I describes all antibodies that link to the RBD in the *up* configuration. This antibody corresponds to the gene *VH3-53* and blocks the binding to the ACE2 receptors. These include antibodies named B38, C102, C105, (*17*–*19*). Class II groups all antibodies that link to the RBD motif blocking the binding with the ACE2 receptor in the *up* and *down* configurations. Class II hNAbs includes highly effective hNAbs named C002, C119, C144 and P2B-2F6 (*17, 20*) corresponding to the gene *VH3-53 and VH4-38*02* with longer complementarity-determining regions than Class I. Class III refers to hNAbs that bind *outside* the RBD motif and bind to both the *up* and *down* configurations. This class of antibodies shows remarkable binding properties that are distinct from the other classes. Class III antibodies includes C135, C110, S309, and REGN10987 (*17, 21, 22*). Barnes *et al.* (*17*) classified Class IV as involving previously described antibodies that do not block the ACE2 and bind only *up* RBDs. Unfortunately, Class IV antibodies did not show much promise to block COVID19 infections. An additional class of hNAbs against SARS-CoV-2 has been identified, particularly antibodies that bind *outside* the RBD motif in the S-protein. For instance, Chi *et al.* (*23*) have shown that a potent antibody links to the N-terminal domain of the S-protein of SARS-CoV-2. We shall refer to this type of antibody as Class V. Here, we focus our attention on Class I, II, and III hNAbs.

Despite detailed information about these antibodies’ structural and biophysical aspects, essential questions remain unanswered. For instance, the discovery of rapidly spreading new SARS-CoV-2 lineages with mutations in the S-protein, particularly in its RBD, has raised concerns among the health authorities and scientific community. It remains unclear whether hNAbs developed by patients during infections to viruses containing the original S-protein sequence will remain effective to these new strains. Moreover, it is also unknown if the new strains propagate faster due to an easier attachment to the receptors or other biological factors. Finally, the different hNAbs classes’ detailed binding mechanisms remain unavailable (*16, 24*–*26*). This information is vital because it can enable researchers to understand what type of antibodies are more efficacious to bind to the S-protein and aid effective therapeutics with antibodies against COVID19. Molecular simulations can be used to answer some of these questions and identify key molecular binding mechanisms between SARS-CoV-2’s new strains and hNAbs. This information can be used to understand how viral mutations impact vaccine developed immunity and to generate treatments that are robust against virus evolution (*16, 24*).

This work investigates the mechanisms for binding and strategies to escape several hNABs used by SARS-CoV-2 strains using molecular simulations. We pursued the same approach as described by Ponga (*6*) to compute binding energies and detailed in **Methods**. Remarkably, this approach predicted the binding affinity and detachment force between the S-protein and ACE2 receptor in good agreement with experimental measures (*1, 3, 5*). We benchmarked five hNAbs (two in Classes I and II and one in Class III) with the RBD, including three mutations N501Y (MT1), E484K, and N501Y (MT2), and K417N, E484K, and N501Y (MT3). These mutations resemble similarities with the novel strains B1.1.7 (*27, 28*), B1.1.28 (*29*), and B1.135 (*30*), respectively. A summary of the simulations can be found in Table 1.

**Table 1.** Details of the simulations used in this work, including the antibody name and PDB file used in the simulations. Total number of atoms in the simulation includes the proteins, and solvent used in the different simulations. Number of Cl^−^ ions used to balance the charge in the simulations. The Class of the antibodies is also specified. The symbols indicate whether the antibody links to the RBD in the up (↑), up/down (⇅), or up/down from the side (⇅ ←). The last three columns indicate if the antibody was modeled with mutations MT1: N501Y, MT2: E484K, N501Y, and MT3: K417N, E484K, N501Y.

| Antibody | PDB | Atoms | Protein | Solvent | Ions | Class | MT1 | MT2 | MT3 |
| --- | --- | --- | --- | --- | --- | --- | --- | --- | --- |
| B38 ↑ | 7BZ5 | 106957 | 9476 | 97473 | 8 Cl<sup>-</sup> | I | Yes | No | Yes |
| C102 ↑ | 7K8M | 106829 | 9283 | 97539 | 7 Cl<sup>-</sup> | I | Yes | No | Yes |
| P2B-2F6 $\uparrow\downarrow$ | 7BWJ | 106909 | 9476 | 97422 | 11 Cl<sup>-</sup> | II | No | Yes | No |
| C144 $\uparrow\downarrow$ | 7K90 | 106741 | 6393 | 100344 | 4 Cl<sup>-</sup> | II | No | Yes | Yes |
| C135 $\uparrow\downarrow\leftarrow$ | 7K8Z | 106750 | 6259 | 100491 | 3 Cl<sup>-</sup> | III | No | No | No |

Our results indicate that the landscape for molecular bonding and binding between antibodies and RBD is vast, and several mechanisms are available. Of tremendous interest, we found that Class III hNAbs −that link in the *up* and *down* states outside the RBD motif− have the largest binding energy to the wild type (WT) RBD (~ 80 kJ·mol^−1^). Class II antibodies, that link in the *up* and *down* states in the WT RBD motif, followed (~ 75 − 80 kJ·mol^−1^). Class I hNAbs produced smaller binding energies (~ 60 − 70 kJ·mol^−1^). We found that the binding energy levels are well separated between different classes and all hNAbs produced similar or higher binding energies than the RBD/ACE2 receptor (66.9 ± 3.7 kJ·mol^−1^) (*6*).

We also investigated the effect of mutated S-proteins. We found that while mutation N501Y does not enhance the binding energy between RBD and ACE2, mutations MT2 (E484K/N501Y), and MT3 (K417N/E484K/N501Y) do enhance the binding avidity between RBD and ACE2. This result could partially explain the rapid propagation of the B.1.351 strain in different parts of the world. We also investigated the mutations’ effect on the binding affinity to several hN-Abs, including B38, C102 (Class I), P2B-2F6, and C144 (Class II). The impact of the mutations is antibody-dependent. However, in general, we found less affinity between mutated S-protein and these antibodies, except for B38. Of particular interest, Class II antibodies showed a sharp decrease in their binding energy when mutations MT2-3 were simulated. This result indicates a much lower efficiency to stop the viral infection.

## 2 Results

We first focused on understanding the effect of the mutations on the binding to the human ACE2 receptors. Fig 1(a) shows the binding energy for the WT, as well as for the RBD with mutations MT1, MT2, and MT3. We observed that the WT’s binding energy is in the neighborhood of 70.6 ± 3.7 kJ·mol^−1^. At the same time, MT1 has slightly smaller binding energy values, e.g., 59 ± 3.3. On the other hand, we observed higher binding energy values for MT2 and MT3, reaching about 73.3 ± 3 kJ·mol^−1^, and 83.0 ± 4 kJ·mol^−1^, respectively. We also noted that the binding energy between RBD and ACE2 receptor is close to reported values reported in literature (*6*). The slight difference could be due to differences in the geometry of the RBD. The PMF evolution between RBD and ACE2 for WT and MT1-3 can be found in **Fig. S1**(a).

**Figure 1:**
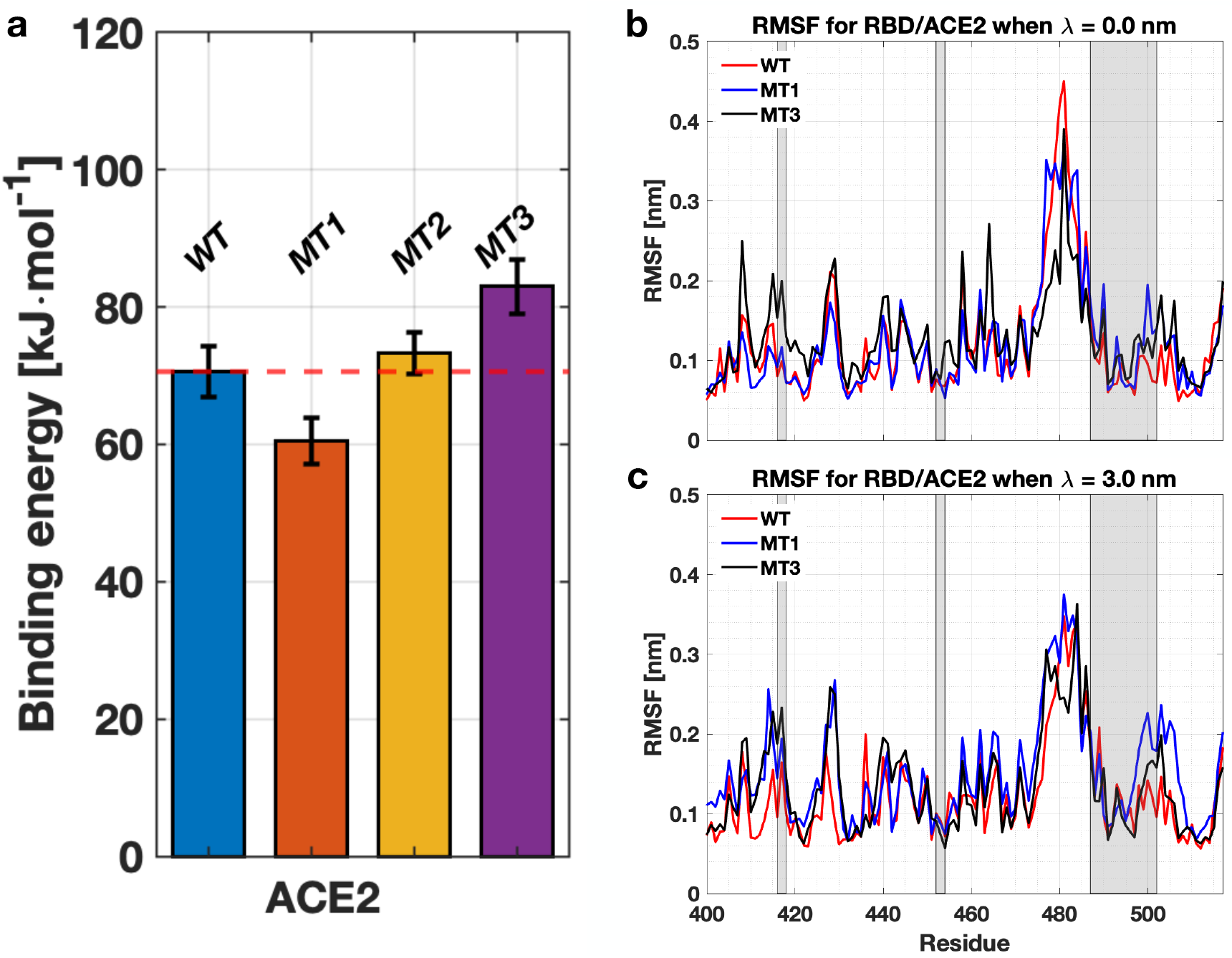
Binding energy between the RBD and the ACE2, for the wild type (WT), mutation MT1: N501Y, MT2: E484K/N501Y, and MT3: K417N/E484K/N501Y. (b) RMSF for residues #400 517 in the RBD for the original and mutated versions (MT1 and MT3) of the RBD in the equilibrium configuration (λ = 0.0 nm). (c) RMSF for the same residues as in (b) when the two particles are detached (λ = 3.0 nm). The shaded areas represent the residues that are in contact with ACE2. RMSF for MT2 is omitted due to overlap with MT3.

In order to obtain a better understanding of the effect of the mutation in the RBD, we computed the root mean square fluctuation (RMSF) for the WT and mutated versions for the equilibrium configuration (λ = 0 nm), and when the proteins where completely detached (λ = 3 nm). The RMSF is shown in Fig. 1(b) and (c) for λ = 0 and 3 nm, respectively. The shaded zones represent residues in contact with ACE2. We observed that there were only small differences between the two versions of the RBD for most residues due to the stochastic nature of the MD simulations. However, larger differences appear in the RMSF near residue #501, both in the equilibrium and detached configurations as shown in Fig. 1(c). Results for MT2 are omitted due to similarities with MT3.

The higher binding energy between RBD containing MT2 and MT3 can be explained by analyzing the residues in contact. Besides having fewer contacts, MT2 and MT3 have a considerably larger binding energy than WT and MT1. We observed the introduction of K417N eliminates the salt bridge between RBD (K417) and ACE2 (D30). The introduction of E484K generates another one between K484 and E35 in ACE2. When looked carefully, we observed that the distances between the new bridge are much smaller (~ 0.13 nm for MT3 compared with ~ 0.3 nm in the MT1, see **Tables S1-3**). The significantly lower distance makes stronger bonds, explaining the higher binding affinity between MT2-3 and ACE2.

### Binding energy between RBD and hNAbs

Next, we investigate the binding energy of five hNAbs described above with the SARS-CoV-2 RBD. The WT S-protein and the three mutations MT1, 2, and 3 were included in the analysis. Let us first describe the residues in contact between the WT S-protein and the hNAbs shown in Fig. 2(a). The red letters correspond to the amino acid residues in contact with the different hNAbs. The underlined letters show the positions of the mutation. Fig. 2(b) shows the location of these residues in the RBD for each antibody with the locations of the selected mutations placed on the RBD. We observed that the mutations were placed on strategic locations of the S-sequence, with the only unaffected binding site unaltered for C135. In particular, K417 generates interactions with three antibodies (B38, C102, C144), E484 interacts with two antibodies in Class II (P2B-2F6 and C144). In contrast, N501 interacts with Class I hNAbs (B38 and C102). The analysis allowed us to strategically made simulations with the mutated RBD with antibodies as detailed in Table 1

**Figure 2:**
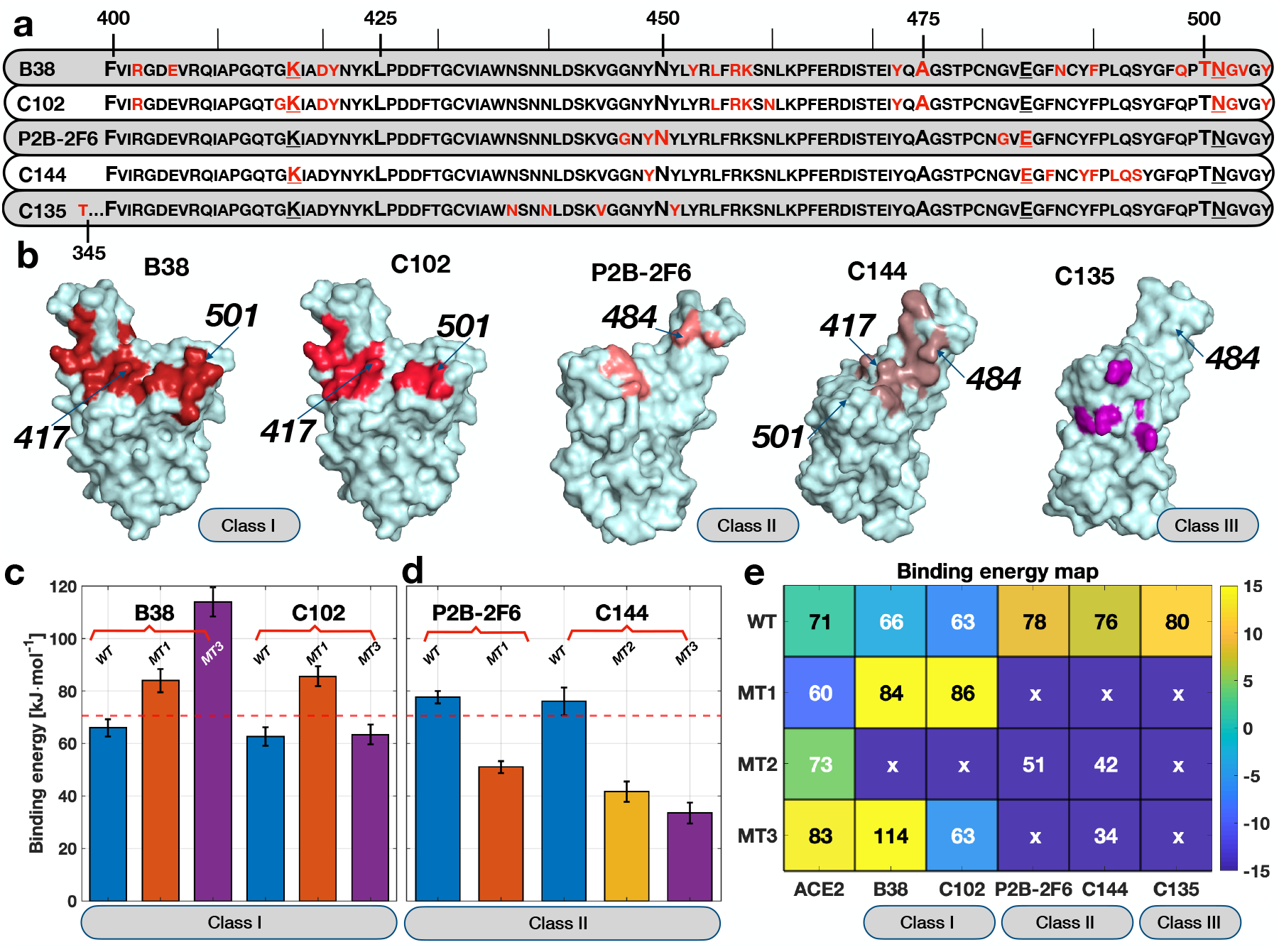
(a) RBD one letter sequence with contacting residues shown in red for the different hNAbs studied in this work. Amino acid residues in the WT sequence that were mutated are shown with an *underline* and larger font size. (b) Location of the amino acid residues in the RBD that interact with the different hNABs. The location of the simulated mutations is also shown in the RBD geometry shown as a surface. (c) Binding energy values between the RBD (WT and mutated) and Class I hNAbs. These mutations include MT1: N501Y, MT2: E484K/N501Y, and MT3: K417N/E484K/N501Y. (d) Binding energy between the RBD (WT and mutated) and Class II hNAbs. (e) Binding energy map between RBD, ACE2, and several hNAbs. The color map indicates the difference in binding taking as a reference the WT RBD and the ACE2 receptor. The number in each cell is the actual binding energy in kJ mol^−1^. The ‘x’ symbol indicates that the binding energy was not computed for that combination since the mutation did not affect the binding residues.

Let us now focus on the analysis of the binding energy between Class I hNAbs and the RBD. As shown in Fig. 2(c), the binding energy between the WT and B38 increased to ~ 66 ± 3.3 kJ·mol^−1^ when all contacts were lost at approximately λ = 1.5 nm (see **Fig S1(b)**). For MT1 and MT3 the last contacts were lost around λ = 2.0 nm reaching a maximum PMF value of ~ 84 ± 4.4, and ~ 114 ± 4.4 kJ·mol^−1^, respectively. The significant difference in the binding energy between mutated and original RBD is explained in the next section’s residues interactions.

Let us now analyze the binding energy for Class I hNAb C102, shown in Fig. 2(c). Remarkably, the binding energy between the WT RBD was in the neighborhood of ~ 62.7 ± 3.55 kJ·mol^−1^, while for MT1 and MT3 it reached ~ 85.6 ± 3.8, and ~ 63.4 ± 3.8 kJ·mol^−1^, respectively (see Fig 2(b) for position of the mutations). The similar binding energy between ACE2 and C102 indicates a similar dissociation constant, illustrating direct competition between receptor and antibody for linking to the S-protein. Like B38, MT1 increased the binding energy for C102. However, the combination in MT3 significantly reduces the binding energy to levels below those found for the WT and ACE2. Thus, while C102 remains active under MT1 (e.g., B.1.1.7 strain), it might lose efficiency against MT3 (e.g., B.1.351 strain). An evolution of the PMF for C102 is shown in **Fig S1(c)**.

We now proceed to quantify the binding energy for Class II hNAbs, in particular, for P2B-2F6 and C144 hNAbs. The binding energies and PMF are shown in Figs. 2(d) and **Fig. S1(d)-(e)**, respectively. The binding energy between the WT S-protein and P2B-2F6 was measured to be ~ 77.7 ± 2.4 kJ·mol^−1^. The significantly higher binding energy compared with the ACE2 receptor indicates a strong binding configuration −as denoted in the PMF plot shown in **Fig. S1(d)**− and illustrates why P2B-2F6 is a potent hNAb. On the other hand, we observed that MT2 −which includes mutation E484K− has a deleterious effect on the binding energy for P2B-2F6, reducing it to ~ 51 ± 2.3 kJ·mol^−1^. This reduction of the binding energy −due to removing a salt bridge between RBD and P2B-2F6 (see next section)− renders the antibody much less effective to bind to the S-protein.

For C144, another Class II hNAb, we found a similar behavior. The binding energy between WT RBD and C144 was ~ 76.1 ± 5.25 kJ·mol^−1^, and the PMF (see Fig. 2(d) **Fig. S1(e)**) was characterized by a strong binding configuration. The similar binding level between WT and Class II hNAbs is remarkable. It illustrates why these types of antibodies are effective against WT SARS-CoV-2. However, the impact of the mutations is drastic for C144. We observed that both MT2 and MT3 produce harmful effects on the binding of the antibody and S-protein. The binding energy was reduced to minimum levels of 41 ± 3.9 and 33.5 ± 3.9 kJ·mol^−1^ for MT2 and MT3, respectively. Unfortunately, the severe reduction in the binding energy suggests that C144 will not be effective against the S-protein’s mutated version (e.g., against the B.1.351 strain).

Next, we analyze the behavior for C135 hNAb that attaches outside the RBD motif. The total binding energy for C135 is close to ~ 80.5 ± 5 kJ·mol^−1^. In particular, the simulation setup was such that the S-protein slid over C135, generating a shearing mechanism. As a result, we found a well-defined energy indicating a stable binding configuration. The PMF increased almost monotonically with λ until the total detachment was performed (see **Fig. S1(f)**). This notable binding energy illustrates the power of Class III hNAbs to block the S-protein. Since the residues in contact between RBD and C135 did not involve any of the mutated versions, we omitted the analysis of MT1-3 (see Fig 2(b)). Fig. 2(e) shows a binding energy map summarizing all binding energies calculated in this work. Cells with ‘x’ indicated that the particular combination of RBD and antibody was not simulated since the mutated residues did not interact with the antibody. The color scale in the map is the binding energy difference, taking the RBD/ACE2 receptor as a reference. The actual values are shown in each cell, with white fonts for those combinations where the binding energy is below the reference and with black fonts for those combinations above the reference.

### Mechanisms for molecular binding of hNAbs

It is now beneficial to compare the molecular mechanisms for binding between the S-protein and hNABs. Fig. 3 shows the antibodies Fabs linked to the RBD, with several insets showing the residues in contact due to Hydrogen (H) bonds and salt bridges. The contacts are shown for locations near the mutation points discussed above.

**Figure 3:**
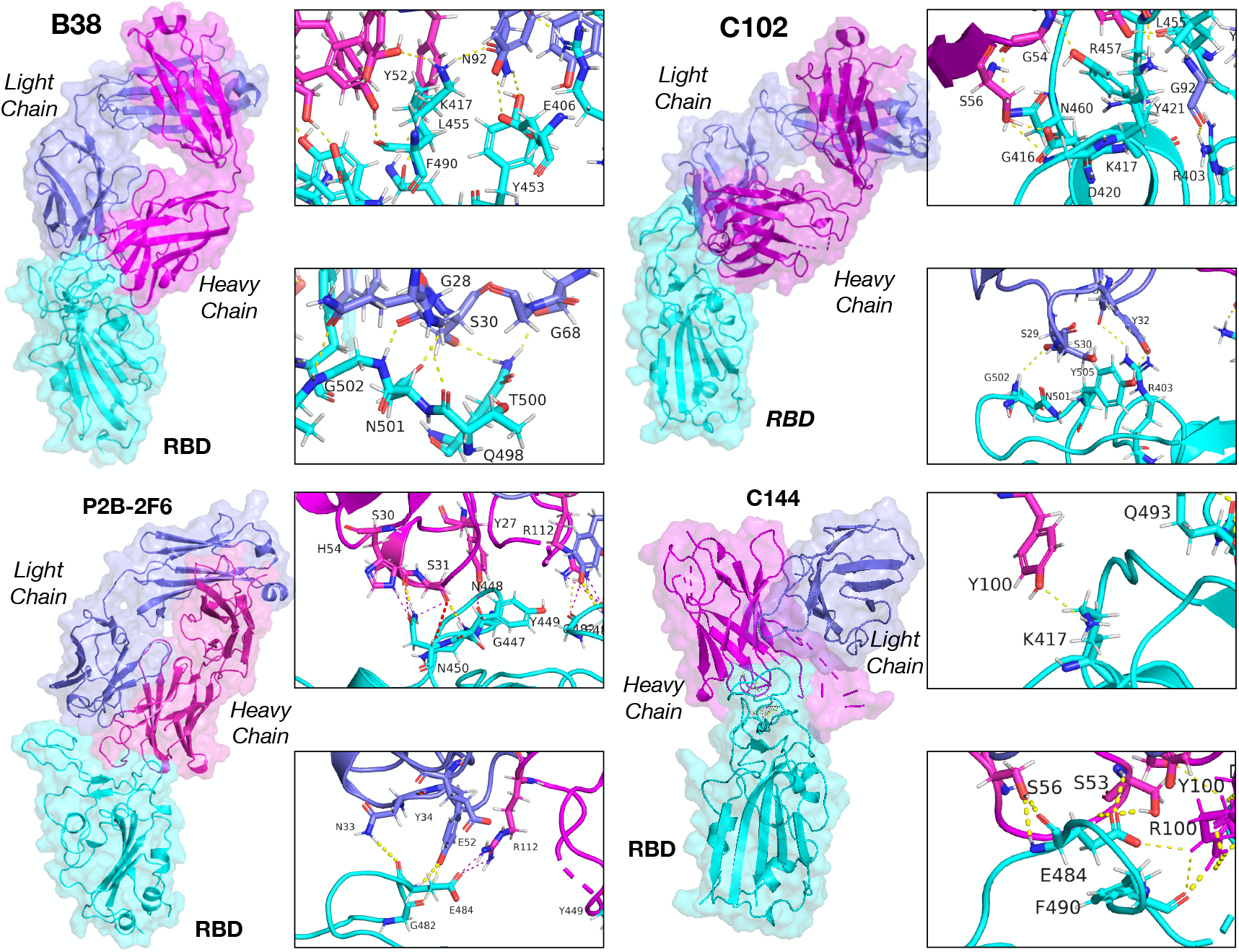
Antibodies Fabs link to the S-protein RBD structure. The RBD is shown in cyan while the HC and LC are shown in magenta and blue, respectively. (a) Structure of B38 and RBD, with the insets show a detailed view of the residues with H-bonds near K417 and N501. b) Configuration for C102 with residues showing H-bond in the insets near K417 and N501. c) P2B-2F6 linked to the RBD with residues in contact near N448 and E484. d) C144 linked to the RBD with residues near K417 and E484.

#### Class I antibodies

**Table S4** shows the amino acid residues in contact between the hNAb B38 and the RBD for the equilibrium configuration (λ = 0.1 nm), including a detailed description between the residues in the heavy chain (HC) and light chain (LC), as well as the ones in the RBD with the distances between these residues. For λ = 0.02 − 0.13 nm, we observed that all twenty-five residue interactions were present via H-bonds without salt bridges. Thirteen of these interactions were made by the HC, while the LC made twelve. We observed that K417 interacts with the HC and LC, with residues Y52 and N92. Residue N501 interacted with residue S30 in the LC with a distance of ~ 0.27 nm (see inset in Fig. 3(a)). As the reaction coordinate increased, the number of residues in contact diminished as shown in **Tables S5-7**. Before complete detachment, at λ = 1.3 nm, residue N487 in the RBD was in contact with S31 in the HC. The analysis indicates a stronger interaction of the RBD with the HC for B38.

Besides using the same geometry for MT1, we observed (see **Table S8-11**) that the number of contacts was lower for the mutated RBD with B38. However, the HC increased the number of contacts (sixteen), while the LC decreased them (five). It is this shift in the contacts that is responsible for the higher PMF. Of remarkable interest, residue Y501 does not play a role in the contact with the B38 in the equilibrium configuration (see Fig. 3(b)), but it does interact with S30 in the LC when λ = 0.2 − 0.5 nm (see **Table S10**). For the MT3 mutations, we observed that the mutated residues did not interact in the equilibrium position between B38 and the RBD (see **Table S12**).

The interacting residues for the hNAb C102 are detailed in **Table S13** for the equilibrium configuration (λ ~ 0.1 nm). The similarities between B38 and C102 are quite remarkable. Out of the fourteen residues in the RBD contacting C102, eleven were shared with B38, and all interactions were due to H-bonds (see **Table S13**). The lesser number of contacts found in C102 explains the lower binding energy for C102 compared with B38. A key difference is that as the reaction coordinate increased, the LC lost contact with the RBD (see **Table S14**). Of tremendous interest, one of the last residues in contact was between N487 in the RBD and with S53 in the heavy chain, which is very similar to the last contact for B38 (see **Table S15**). This observation suggests that the engagement between RBD and the hNAbs could start in residue N487 of the RBD.

The contacts between MT1 RBD and the hNAb C102 are detailed in **Table S16-19** for several values of λ. The HC of C102 is the main responsible for the contacts (ten out of thirteen). However, the LC’s contacts were maintained until λ ~ 0.55 nm, justifying the higher PMF observed in **Fig. S1(c)** in comparison with the original RBD. In the analysis, we did not find evidence that Y501 interacted with the antibody, unlike the original RBD. For MT3, we observed that residue N417 interacted with the HC. However, the other mutations did not interact with the antibody (see **Table S20**).

#### Class II antibodies

Let us now focus on hNAbs Class II, namely P2B-2F6 and C144, whose interacting residues with the RBD are detailed in **Tables S21-22**. We observed a different binding strategy than Class I hNAbs, with fewer contacts but with a much stronger binding energy. For the hNAb P2B-2F6, we identified five residues interacting in the RBD with seven residues in the antibody (five in the HC −Y27, S31, S30, H54, and R112− and two −Y34 and N33− in the LC). A key difference in the binding strategy is that in addition to the H-bonds, we observed a salt bridge between R112 and E484 (see inset in Fig. 3(c) and **Table S21**). As λ increased, the number of residues decreased. Y43 and R112 in the P2B-2F6 and E484 were the last bonds to be broken, indicating the large interaction forces between these two residues.

For C144, a similar binding strategy was observed. Seven residues in the RBD were in contact with eight in the HC hNAb. The LC did not interact with the RBD since the HC makes direct physical contact with the RBD (see Fig. 3(d)). Another salt bridge between E484 and R100 in the HC was observed (inset in Fig. 3(d)). The simulation of Class II antibodies illustrates the role of salt bridges and how they can serve to initiate the binding between the two particles.

Mutations MT2-3, involving a change in E484K, explain this mutation’s deleterious effect in the binding between RBD and P2B-2F6 and C144. The direct removal of the salt bridge reduces the binding energy significantly for both cases. While K484 interacted with the HC, it was through a weak H-bond with S55 (r = 0.26 nm). Also, nine residues in the RBD were in contact compared to eleven in the WT, explaining the lower binding energy.

#### Class III antibodies

We now describe the bonding between C135 Class III hNAbs and the RBD. Fig. 2(e) shows the remarkable differences between Class III hNAbs with the other classes. We observe that Class III targets residues T345 and residues between N437-Y451 as illustrated in Fig. 2(e). We also observed no overlap between residues in Class III and Class I - II, suggesting that combined strategies for neutralization are possible. C135 uses seven residues (five in the HC and two in the LC) to bind to five residues in the RBD (see **Table S24** and inset in Fig. 4). All these bonds were H-bonds, without any salt bridges. However, since the mechanism for detaching is due to sliding on the surfaces of the RBD, it generated long-lasting contacts (and even new ones −e.g., G97 with T345 and R346 with E50− as shown in **Table S24**). Thus, as the two molecules were pulled apart, the sliding mechanism led to increased binding energy compared to other hNAbs classes.

**Figure 4:**
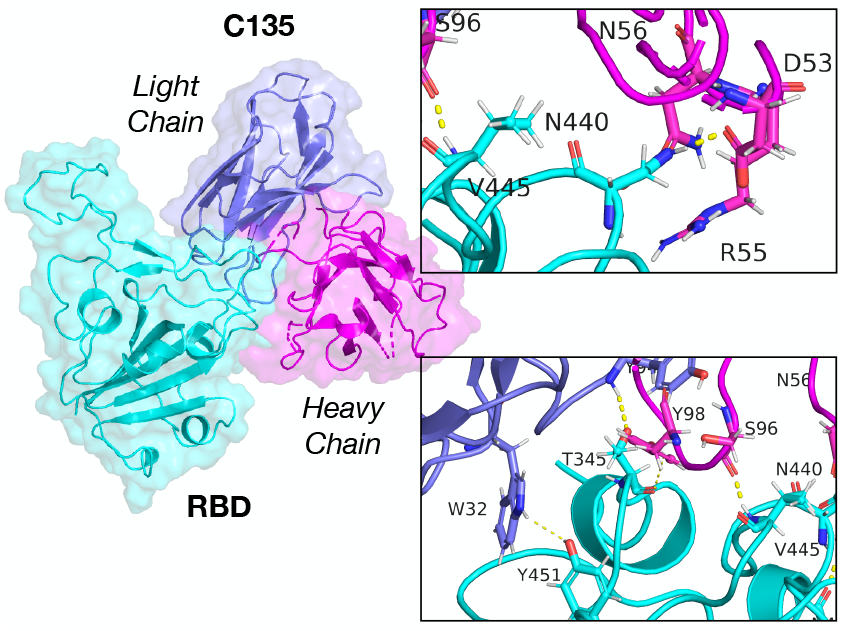
C135 antibodies Fabs link to the S-protein RBD structure. The RBD is shown in cyan while the heavy chain (HC) and light chain (LC) are shown in magenta and blue, respectively.

## 3 Discussions

We have investigated the mechanisms and strategies for antibody binding and escape in SARS-CoV-2. On the one hand, analysis of the WT and MT1 of the RBD and the ACE2 receptor indicates that the binding energy remains at similar levels and between our simulations’ margin error. On the other hand, for MT2 and MT3 RBD versions involving mutations K417N/E484K/N501Y, the binding energy increased compared to the WT. This observation suggests that the strain B.1.351 could have a better binding avidity with the ACE2 receptor and better effectivity to enter host cells than the WT virus. This observation seems to agree with recent works where the effect of the mutations have been analyzed. In particular, several experimental works using mutated RBD have shown a more pronounced effect of the mutation E484 (*16, 25, 31–34*). We also observed that the mutations increased the RMSF near the mutation zone, suggesting more residues’ motion to bind with the receptor. The larger values of the RMSF indicate contacts that are on and off at the interface in the mutated versions. However, the biological effects on the overall higher RMFS near residue #501 are unclear since larger fluctuations leads to more exposure of the residues, which in turn could lead to a more efficient attachment (increase in the probability of attachment).

The binding mechanisms between RBD and hNAbs are vast, and various classes of antibodies use different strategies. Based on our study, Class II and III hNAbs are the most effective to bind to the WT RBD. Considering that Class II and III link to different RBD areas, therapeutics strategies combining Class II and III could be very effective to stop viral infection as suggested by others (*17*). Unfortunately, we observed that Class II hNAbs show a sharp decline in the binding energy to MT2 and MT3 and are due to two main factors. First, the loss of the salt bridge made by residue E484 and Class II hNABs that use gene *VH3-53* as a germline (and the same strategy has been seen for P2B-2F6 that is different gene). Second, the reduced number of contacts between Class II and RBD (see **Tables S21-23**) makes the binding energy decay to low levels. This renders Class II antibodies inefficient to bind to the RBD. This observation suggests that Class II hNAbs are less effective in binding to the S-proteins within the B.1.351 strain (*16,25,33,34*). The results are in agreement with experimental observations. For instance, recent experimental work on mutated RBD with E484K has shown a 100-1000 fold increase in antibody dose for C144 (*16, 25*). This effect is more drastic in C144 since it uses the HC to make contact with the RBD, while P2B-2F6 uses both the HC and LC (see Fig. 3(c)-(d)).

Simulations between the RBD and Class I hNAbs indicated that the binding energy increases between MT1 (N501Y) and B38 and C102 compared to WT. This observation suggests that Class I hNAbs will remain effective to bind to the S-protein for the B.1.1.7 strain. This result is supported by experimental works where N501Y mutation has been found to have a modest effect on the binding of monoclonal antibodies (*15, 25*). The binding energy between B38 and C102 had different behavior when analyzed with MT3. B38 had shown higher binding energy 114 kJ·mol^−1^. On the other hand, C102 showed binding energy significantly below the value found for the ACE2 receptor and MT3, indicating less effectivity to bind to the mutated S-protein. Once again, this observation suggests that C102 will be less effective in blocking viral infection for viruses with the B.1.351 lineage.

Our all-atom MD simulations also have several limitations. For instance, to reduce the computational cost, only part of the S-protein was modeled in this work. Previous studies have shown minor differences in the PMF plot for one single RBD and the full trimeric S-protein (*6*). However, the complete simulation of the S-protein with glycans can elucidate additional binding mechanisms with antibodies. Additionally, the mutated RBD has been modeled, but cryoEM mutated molecular structures can differ from those used in this work. As far as the authors are concerned, there is not cryoEM of mutated RBD available yet. Differences in the geometry could affect the equilibrium position and, thus, change the energy path. While these limitations can change the actual values found in this work, we argue that the trends will remain the same as the molecular mechanisms will remain mostly unaffected.

In conclusion, we have investigated the mechanisms and strategies for antibody binding and escape in SARS-CoV-2. Analysis of the WT and several mutated versions of the RBD and the ACE2 receptor indicates that the binding energy remains at similar levels as the WT for N501Y but increases for mutations K417N and E484K. This observation suggests that strain B.1.351 could have a better dissociation constant and a better effectivity to enter host cells than the WT virus. The effect of the mutations in antibodies’ avidity of these mutations is mainly detrimental, and most antibodies reduce the binding energy in the presence of mutations, except B38. Future directions of this work include the computational modeling of hNABs with better binding affinity to the mutated RBD using data-based approaches and the estimation of antibody dose using mechanistic models.

## Materials and Methods

### Molecular dynamics simulations, Umbrella sampling and characterization

Molecular dynamics simulations were performed with GROMACS (*35*–*37*) simulation package. Molecular geometries were obtained from different protein database (PDB) files, including the 6LZG (RBD and ACE2) (*38*), 7BWJ (P2B-2F6) (*20*), 7BZ5 (B38) (*19*), 7K8M (C102), 7K8Z (C135), 7K90 (C144) (*17, 18*). For the antibodies given in files 7BWJ, 7BZ5, 7K8M, the database’s molecular geometry includes the SARS-CoV-2 geometry of the RBD in solution with the antibodies. For 7K8Z, 7K90, the PDB files included the full trimeric proteins of the SARS-CoV-2. To make fair comparisons and reduce the simulation runtime, we used a portion of the S-glycoprotein, limited to the RBD between residues 337−505 (NCBI Reference: YP 009724390.1). Since some of the PDB files’ sequences were incomplete, we used an RBD from 7K8M and aligned the proteins using Pymol. The use of the same RBD ensures the same geometry for all antibodies with a root mean square displacement (RMSD) per atom less than 0.075 nm.

The antibody geometries with the RBD were processed in GROMACS, where the proteins’ principal axis was aligned along the x−direction, which was then used as the main reaction coordinate for the steered MD simulation. Some of the protein files had missing atoms that were not resolved in the cryoEM identification. We included these atoms using the SwissViewer. Once the proteins were complete, we solvated them in water using the TIP3P model to achieve a density of approximately *ρ* = 1000 Kg·m^−3^. In order to allow for sufficient space for the steered simulations, we generated computational cells with more than 1 nm between the proteins and end of the cells and sufficient space on top to perform the pulling simulations. The computational cell had dimensions of ~ 20 × 7.55 × 7.25 (nm). After adding the solvent, the system had a non-zero charge, and ions (Cl^−^) were added as needed to equilibrate in all samples.

All interatomic forces were computed with the CHARMM force-field (*39*). The computational cells had around ~106,000 atoms, including proteins and solvent. The details of the antibody simulations is shown in Table 1.

After solvating the system with water, we minimized the system’s energy using a non-linear conjugate gradient, with a force tolerance of 1000 kJ·mol^−1^·nm^−1^. The relaxed cells were then subjected to a constant temperature (*T* = 310.15 K), and pressure (*P* = 1 bar) run for about *t* = 2 ns using a Berendsen thermostat (*40*). A timestep of Δ*t* = 2 fs was used for all simulations. Short-range interactions were treated with a smooth force-switch cutoff of *r* = 1.2 nm, and long-range electrostatics were treated using the Particle-Mesh-Ewald (PME) formalism, implemented in GROMACS (*41*) for all simulations. Hydrogen−bonds were restrained with the LINCS algorithm (*42*).

To compute the binding affinity between the RBD of the S-proteins and several antibodies, we used a combination of steered MD simulations with umbrella sampling. In all simulations, the antibody (Heavy and Light) chains were fixed in the simulation cell using a position restrain, while the RBD was subject to a pulling force acting on the x– direction. This force was applied using a spring constant of *K* = 1000 kJ·mol^−1^·nm^−2^. The force was applied between the center of mass between the RBD and the antibody. The proteins were then pulled apart from each other at a constant speed of *v_x_* = 1 nm·ns^−1^ along the x-direction.

Configurations were obtained every 20 ps in order to sample the PMF along with the reaction coordinate. We selected configurations separated by a distance of Δλ = 0.05 nm or less in most cases. These configurations were then relaxed using a short NPT ensemble run of 0.1 ns for a posterior sampling of the PMF using the umbrella technique. Umbrella simulations were performed using NPT ensembles for about 10 ns, using a spring constant of *K* = 1000 kJ·mol^−1^·nm^−2^ between the center of mass of the molecules. The PMF was computed with the Weighted Histogram Analysis Method (WHAM) (*43*) using a bootstrap analysis to estimate the uncertainty in the PMF. We used 200 different measures, using 200 binning spaces along with the reaction coordinate λ and a tolerance of 10^−8^. Instead of using the moment when the two molecules detach physically as Ponga (*6*), the binding energy was taken as the value when the PMF plateaus, taking the PMF’s average during the last 0.5 nm of the reaction coordinate.

Analysis of the atomistic configurations was performed with the GROMACS cluster analysis tool. We scanned the configurations with a root mean squared displacement between the range of 0.15 − 0.25 nm (*44*). The cluster analysis yielded between three to six clusters for the analyzed configurations. In all cases shown, the most populated cluster was used when analyzing the configurations. The configurations and H-bonds were then analyzed with the software Pymol, and the salt bridges with ESBRI (*45*).

### Mutations of the RBD

To assess the ability of antibodies to link to mutated S-proteins, we used the recently reported mutation of the S-protein corresponding to MT1: N501Y, MT2: E484K/N501Y, and MT3: K417N/E484K/N501Y. The mutations were done manually using the atomic positions given for the RBD in complex modifying the amino acid residues. We selected the rotamer that produced better higher mutation probability and lesser overlap with the protein chain. The configuration was then deformed slightly to remove overlaps (using the clean command in Pymol) within a neighborhood of 0.5 nm of the mutated residues. All samples were then thermally stabilized following the procedure previously described.

## Acknowledgements

We gratefully acknowledge the support from the Natural Sciences and Engineering Research Council of Canada (NSERC) through the Discovery Grant under Award Application Number RGPIN-2016-06114, the New Frontier in Research Fund Exploration program through the award NFRFE-2019-01095 and the support of Compute Canada and the Advanced Research Computing (ARC) at the University of British Columbia.

## Declaration of contribution

M.P. designed the research plan, performed the simulations and analysis. M.H and S. K. performed the analysis of the residues, H-bonds, and salt bridges. All authors wrote and made comments to the manuscript.

## Data and materials availability

All data needed to evaluate the conclusions in the paper are present in the paper and/or the Supplementary Materials.

