## Supplementary Information file for "Molecular strategies for antibody binding and escape of SARS-CoV-2 and its mutations"

March 4, 2021

### 1 PMF as a function of the reaction coordinate for ACE2 and hNABs

**Fig. S1** shows the potential of mean force (PMF) for receptor binding domain (RBD) and the ACE2 receptor and several human neutralizing antibodies (hNABs) as a function of the separation distance, $\lambda$. The shaded area represents the error estimation in the measure obtained from 200 estimations of the PMF.

The evolution of the PMF for the RBD and ACE2 receptor is found in **Fig. S1**(a). We observe how MT2 and MT3 have higher binding energy compared to the WT. For MT1, we found the binding energy to be below the WT.

The PMF evolution as a function of the reaction coordinate for B38 is illustrated in **Fig. S1**(b), including the RBD with the WT, MT1, and MT3. We observed a well-defined equilibrium configuration for B38 −for the original and mutated RBD− with a global minimum at $\lambda = 0.1$ nm. The energy increased almost quadratically up to $\lambda = 0.2$ nm. After that, the PMF increased with irregular fluctuations due to the loss of contacts between the two molecules. For the mutated version, the PMF shows another local minimum around $\lambda = 0.25$ nm due to intermittent contacts on and off. These two minimum configurations are separated by an energy barrier of $\sim 25$ kJ·mol$^{-1}$. The behavior for the MT1 and MT3 versions is then similar, although the actual values differ. The PMF plateaus at around $\lambda = 1.2$ nm except for MT1.

The PMF evolution for C102 is shown in **Fig. S1(c)**, where a non-convex energy landscape with two minima (one global at $\lambda = 0.1$ nm and another local at $\lambda = 0.2$ nm) for both the original RBD and the mutated one can be observed. This behavior indicates multiple equilibrium configurations of the antibody C102 separated by a small barrier of $\sim 10$ kJ·mol$^{-1}$. After that, the PMF increased with the reaction coordinate until the proteins were separated entirely. This separation was manifested in the PMF plot with a plateau at $\lambda = 2.0$ nm and $\lambda = 1.75$ nm for C102 with the RBD in its original and mutated versions (MT1 and MT3).

Next, let us analyze the PMF for Class II P2B-2F6 hNAb, as shown in **Fig. S1(d)**. We found well-defined minimum configurations near $\lambda = 0.1$ nm for both antibodies, denoting a robust binding configuration. The PMF increased sharply for P2B-2F6 as $\lambda$ increased until it reached $\lambda \sim 0.7$ nm. After this point, the PMF plateaus. The behavior for MT2 is similar, although the PMF shows a first plateau region, and then it declines. We have taken the binding energy as the average value of the first plateau.

For C144 hNAb, another one belonging to Class II, the PMF evolution also increased sharply until $\lambda \sim 0.5$ nm (see **Fig. S1(e)**). Then, the PMF showed periods with more or less constant values followed by incremental PMF regions. This behavior indicates sudden detachment of molecules as $\lambda$ increased. Remarkably, when $\lambda \sim 2$ nm, the binding energy plateaued. This observation suggests that Class II has a very well-defined binding energy level, above the levels found for the ACE2 receptor and Class I hNAbs. For MT2 and MT3, we observe that the PMF had a significant lower value. We see that the most significantly decline in the PMF is given for MT2, which involves E484K and N501Y. The decrease in the PMF for MT3 shows the effect of K417N. Nevertheless, these values are well below the binding energy found for C144 and WT RBD.

We finally focus on the PMF for C135. We observed almost a monotonically increasing behavior. The actual physical detachment occurred at $\lambda \sim 2$ nm, where the PMF seemed to have an almost constant value. We have taken the binding energy as the average over that region.

The histograms for PMF sampling between the ACE2 receptor and RBD (WT, MT1, MT2, and MT3) are shown in **Fig. S2**

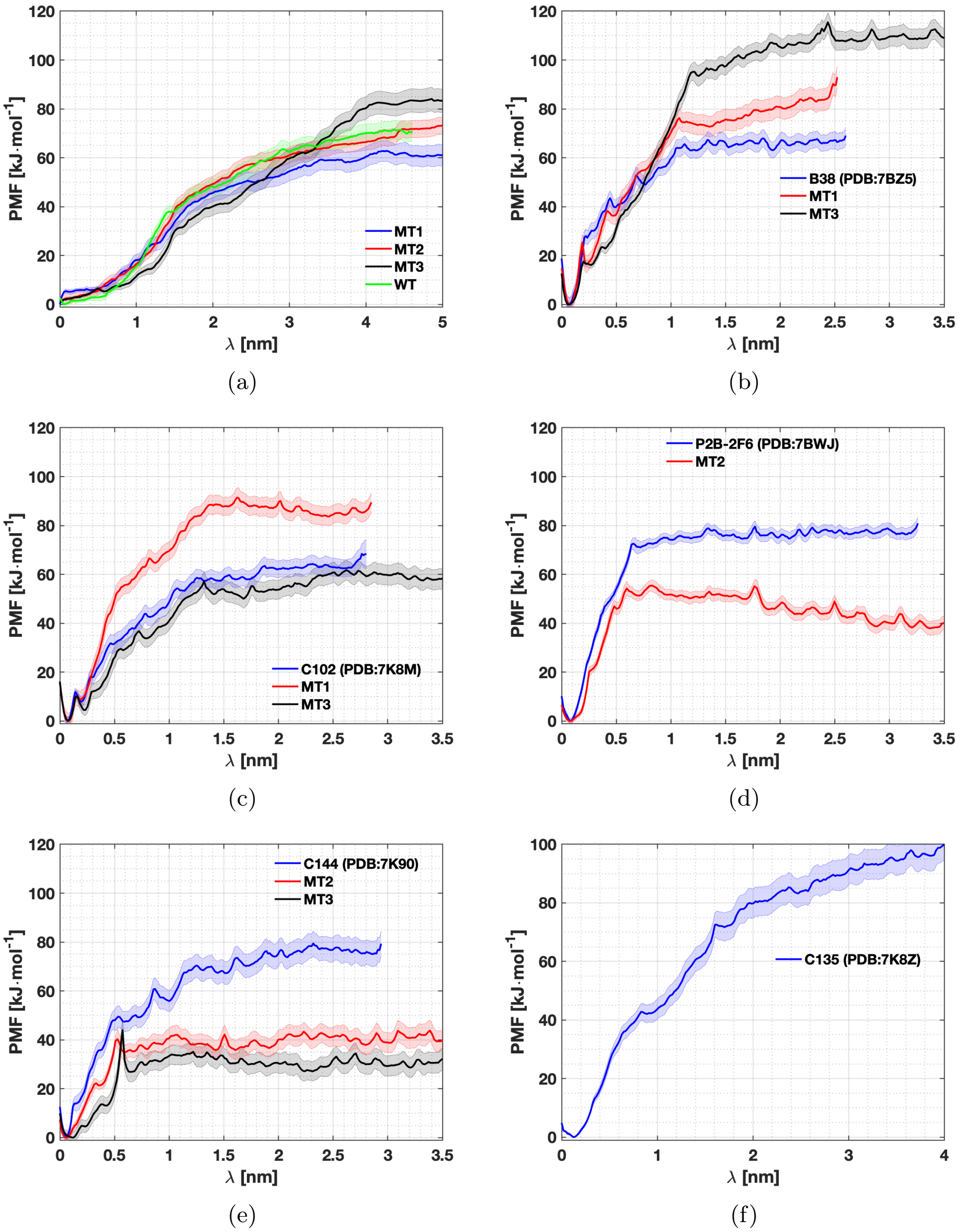


**Figure S 1**: PMF between the RBD and the ACE2 receptor and several hNAbs.

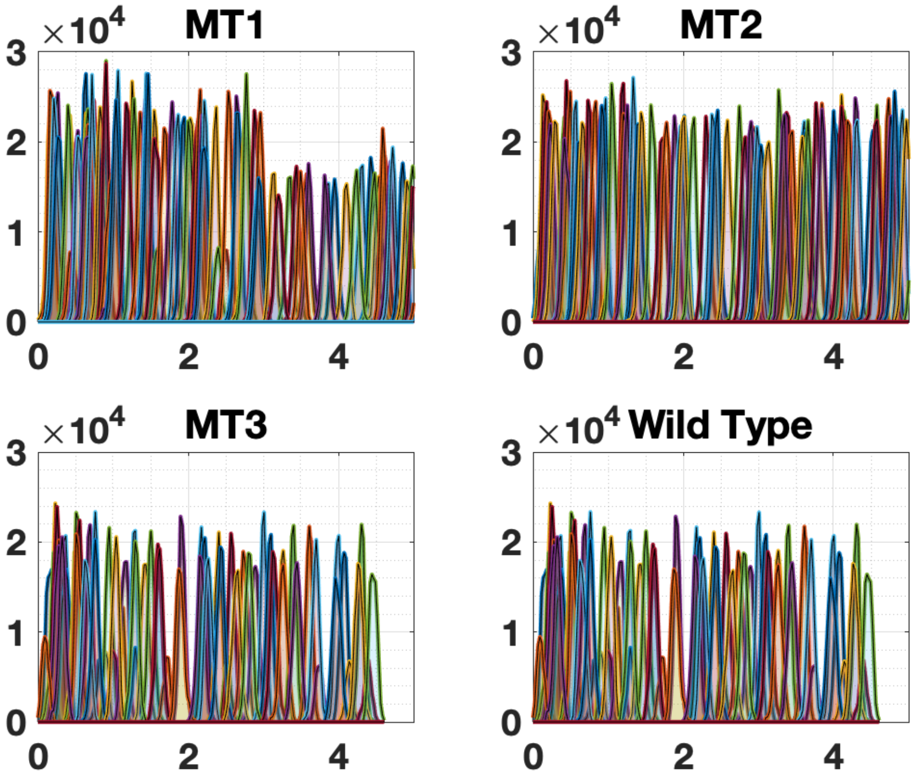


(a)

**Figure S 2**: Histogram showing the PMF sampling of the reaction coordinate for the RBD. The wild type (WT), MT1 (N501Y), MT2 (E484K, and N501Y), and MT3 (K417N, E484K, and N501Y) are shown.

#### 2 Interactions between RBD and ACE2 receptor

Below we present the amino acid residues that are interacting between the RBD MT1 and the ACE2 receptor.

**Table S 1**: The hydrogen bonds and salt-bridges between SARS-CoV-2 RBD containing one mutation (N501Y) and labeled MT1 and the human ACE2 receptor.

| $\lambda = 0 - 0.30$ nm | ACE2 | Length(Å) | SARS-CoV-2 RBD |
|---|---|---|---|
| Hydrogen bond | D30 | 1.8 | K417 |
| | D38 | 1.6 | Y449 |
| | S19 | 1.8 | A475 |
| | S19 | 1.8 | S477 |
| | Y83 | 3.4 | N487 |
| | K31 | 1.9 | Q493 |
| | E35 | 2.3 | Q493 |
| | K353 | 1.9 | G496 |
| | K353 | 1.7 | Q498 |
| | K353 | 1.9 | **Y501** |
| | N330 | 2.0 | **Y501** |
| | D355 | 3.2 | G502 |
| | G354 | 2.9 | V503 |
| | G354 | 1.8 | G504 |
| | E37 | 1.8 | Y505 |
| Salt bridge | D30 | 2.7, 3.9 | K417 |
| $\lambda = 0.27 - 0.60$ nm | | | |
| Hydrogen bond | E23 | 1.8, 2.7 | K458 |
| | Y83 | 3.1 | N487 |
| | Y83 | 1.8 | Y489 |
| | K31 | 2.0 | Q493 |
| | K353 | 2.0 | Y495 |
| | K353 | 2.4 | G496 |
| | Q42 | 2.6 | Q498 |
| | R357 | 2.4, 3.2 | T500 |
| | K353 | 1.8 | **Y501** |
| Salt bridge | E23 | 2.7 | K458 |

**Table S 2**: The hydrogen bonds and salt-bridges between SARS-CoV-2 RBD containing one mutation (N501Y) and labeled MT1 and the human ACE2 receptor.

| $\lambda = 0.62 - 0.85$ nm | ACE2 | Length(Å) | SARS-CoV-2 RBD |
|---|---|---|---|
| Hydrogen bond | D30 | 1.7 | K417 |
| | H34 | 3.2 | Y453 |
| | Y83 | 1.7 | N487 |
| | Y83 | 2.0 | Y489 |
| | H34 | 3.2 | Q493 |
| | D38 | 2.8 | Q493 |
| | K353 | 1.8 | Y495 |
| | K353 | 3.4 | G496 |
| | Y41 | 2.2 | Q498 |
| | R357 | 1.9 | T500 |
| | K353 | 1.7 | **Y501** |
| | D355 | 2.0 | G502 |
| Salt bridge | D30 | 2.6, 3.5 | K417 |
| $\lambda = 1.14 - 1.42$ nm | | | |
| Hydrogen bond | D30 | 2.3 | K417 |
| | Y83 | 1.8 | N487 |
| Salt bridge | D30 | 2.7 | K417 |
| $\lambda = 1.65 - 1.93$ nm | | | |
| Hydrogen bond | Q24 | 2.3 | N487 |
| | Y83 | 1.6, 2.8 | N487 |
| $\lambda = 2.83 - 3.1$ nm | | | |
| Hydrogen bond | S19 | 2.2 | Y473 |
| | Q24 | 2.8 | Y475 |
| | Y83 | 3.4 | N487 |

Below we present the amino acid residues that are interacting between the RBD containing three mutations (K417N/E484K/N501Y) mutation and the human ACE2 receptor.

**Table S 3**: The hydrogen bonds and salt-bridges between SARS-CoV-2 RBD containing three mutations (K417N/E484K/N501Y) and labeled MT3 and the human ACE2 receptor.

| $\lambda = 0 - 0.30$ nm | ACE2 | Length(Å) | SARS-CoV-2 RBD |
|---|---|---|---|
| Hydrogen bond | R357 | 1.8, 3.2 | T500 |
| | K353 | 1.7 | **Y501** |
| | D355 | 2.1 | G502 |
| $\lambda = 0.75 - 1.05$ nm | | | |
| Hydrogen bond | H34 | 2.6 | Y453 |
| | E35 | 1.8 | **K484** |
| | Y83 | 2.1 | N487 |
| | N330 | 3.3 | T500 |
| | D355 | 3.3 | T500 |
| | K353 | 2.6 | **Y501** |
| | K353 | 3.1 | G502 |
| | T324 | 2.7 | V503 |
| Salt bridge | E35 | 1.3 | **K484** |
| $\lambda = 2.25 - 2.60$ nm | | | |
| Hydrogen bond | Y83 | 1.9 | N487 |
| $\lambda = 3.51 - 3.82$ nm | | | |
| Hydrogen bond | E35 | 1.7, 2.5 | **K484** |
| Salt bridge | E35 | 2.1 | **K484** |

### 3 Class I antibodies

Below we present the amino acid residues that are interacting between the RBD (in its WT and mutated versions) and with Class II antibodies B38 and C102.

**Table S 4**: The hydrogen and salt-bridges between SARS-CoV-2 RBD and B38 (PDB:7BZ5) antibody.

| $\lambda = 0.02 - 0.13$ nm | Heavy Chain | Length(Å) | SARS-CoV-2 RBD | Length(Å) | Light Chain |
|---|---|---|---|---|---|
| Hydrogen bond | | | R403 | 1.7 | N92 |
| | | | E406 | 2.9 | Y94 |
| | Y52 | 2.7 | K417 | 2.5 | N92 |
| | S56 | 1.6 | D420 | | |
| | S53 | 3.3 | Y421 | | |
| | G54 | 2.0 | Y421 | | |
| | | | Y453 | 2.4 | N92 |
| | Y33 | 1.7 | L455 | | |
| | S53 | 2.0 | R457 | | |
| | S53 | 2.8 | K458 | | |
| | S31 | 2.0 | Y473 | | |
| | I28 | 2.3 | A475 | | |
| | N32 | 2.2 | A475 | | |
| | G26 | 1.7, 2.8 | N487 | | |
| | R97 | 2.2, 2.5 | N487 | | |
| | Y100 | 3.0, 3.2 | F490 | | |
| | | | Q498 | 2.3 | S30 |
| | | | Q498 | 2.1 | S67 |
| | | | T500 | 3.2 | G28 |
| | | | N501 | 2.7 | S30 |
| | | | G502 | 1.9 | G28 |
| | | | V503 | 3.0 | Q27 |
| | | | Y505 | 2.6 | Q90 |
| | | | Y505 | 2.2 | L91 |

**Table S 5**: The hydrogen and salt-bridges between SARS-CoV-2 RBD and B38 (PDB:7BZ5) antibody.

| $\lambda = 0.06 - 0.19$ nm | Heavy Chain | Length(Å) | SARS-CoV-2 RBD | Length(Å) | Light Chain |
|---|---|---|---|---|---|
| Hydrogen bond | | | R403 | 1.8, 2.9 | N92 |
| | | | Q409 | 3.3 | Y94 |
| | Y58 | 2.7 | T415 | | |
| | | | K417 | 1.8 | N92 |
| | G54 | 3.1 | Y421 | | |
| | Y33 | 1.8 | L455 | | |
| | N32 | 2.3 | A475 | | |
| | G26 | 2.1 | N487 | | |
| | R97 | 2.2, 2.5 | N487 | | |
| | Y100 | 2.9 | F490 | | |
| | Y100 | 2.9 | Q493 | | |
| | | | Y495 | 1.9 | Y32 |
| | | | G496 | 2.0 | S30 |
| | | | Q498 | 2.7 | S67 |
| | | | T500 | 3.1 | G28 |
| | | | G502 | 1.9 | G28 |
| | | | Y505 | 2.0 | Q90 |
| | | | Y505 | 2.5 | L91 |
| | | | Y505 | 2.6 | N92 |

**Table S 6**: The hydrogen and salt-bridges between SARS-CoV-2 RBD and B38 (PDB:7BZ5) antibody.

| $\lambda = 0.09 - 0.31$ nm | Heavy Chain | Length(Å) | SARS-CoV-2 RBD | Length(Å) | Light Chain |
|---|---|---|---|---|---|
| Hydrogen bond | | | R403 | 1.8,2.2 | N92 |
| | | | E406 | 2.8 | N92 |
| | Y52 | 3.4 | K417 | | |
| | G54 | 3.4 | Y421 | | |
| | | | Y453 | 3.1 | N92 |
| | Y33 | 1.7 | L455 | | |
| | S31 | 3.1 | K458 | | |
| | S31 | 2.0 | Y473 | | |
| | I28 | 1.9 | A475 | | |
| | N32 | 2.9 | A475 | | |
| | G26 | 2.9 | N487 | | |
| | R97 | 2.2, 2.9 | N487 | | |
| | R97 | 2.8 | Y489 | | |
| | Y100 | 3.3 | F490 | | |
| | Y100 | 3.0 | Q493 | | |
| | | | S494 | 2.9 | Y32 |
| | | | Y495 | 1.9 | Y32 |
| | | | G496 | 1.9 | S30 |
| | | | Q498 | 2.4 | S30 |
| | | | Q498 | 2.6 | S67 |
| | | | T500 | 3.4 | G28 |
| | | | N501 | 2.9 | S30 |
| | | | G502 | 2.0 | G28 |
| | | | Y505 | 1.9 | Q90 |
| | | | Y505 | 2.3 | L91 |

**Table S 7**: The hydrogen and salt-bridges between SARS-CoV-2 RBD and B38 (PDB:7BZ5) antibody.

| $\lambda = 0.67 - 0.89$ nm | Heavy Chain | Length(Å) | SARS-CoV-2 RBD | Length(Å) | Light Chain |
|---|---|---|---|---|---|
| | N32 | 2.5 | A475 | | |
| | I28 | 1.9 | A475 | | |
| | G26 | 2.3 | N487 | | |
| | R97 | 1.8, 2.5 | N487 | | |
| $\lambda = 0.99 - 1.27$ nm | | | | | |
| Hydrogen bond | S31 | 1.9 | N487 | | |

**Table S 8**: The hydrogen bonds and salt-bridges between SARS-CoV-2 RBD containing one mutation (N501Y) labeled as MT1 in the manuscript and the hNABs B38 (PDB:7BZ5).

| $\lambda = 0.02 - 0.14$ nm | Heavy Chain | Length(Å) | SARS-CoV-2 RBD | Length(Å) | Light Chain |
|---|---|---|---|---|---|
| Hydrogen bond | | | R403 | 1.9, 2.6 | N92 |
| | Y58 | 2.6, 2.8 | T415 | | |
| | S56 | 3.1 | G416 | | |
| | Y52 | 3.3 | K417 | 2.1 | N92 |
| | S56 | 1.6 | D420 | | |
| | S53 | 3.2 | Y421 | | |
| | G54 | 2.1 | Y421 | | |
| | Y33 | 1.6 | L455 | | |
| | S53 | 1.9 | R457 | | |
| | S31 | 3.2 | K458 | | |
| | S31 | 1.9 | Y473 | | |
| | I28 | 2.2 | A475 | | |
| | N32 | 1.9 | A475 | | |
| | G26 | 1.9 | N487 | | |
| | R97 | 1.7, 2.2 | N487 | | |
| | R97 | 3.1 | Y489 | | |
| | Y100 | 3.0, 3.2 | F490 | | |
| | | | G502 | 1.8 | G28 |
| | | | Y505 | 2.5 | Q90 |
| | | | Y505 | 1.9 | L91 |

**Table S 9**: The hydrogen bonds and salt-bridges between SARS-CoV-2 RBD containing one mutation (N501Y) labeled as MT1 in the manuscript and the hNABs B38 (PDB:7BZ5).

| $\lambda = 0.19 - 0.34$ nm | Heavy Chain | Length(Å) | SARS-CoV-2 RBD | Length(Å) | Light Chain |
|---|---|---|---|---|---|
| Hydrogen bond | S56 | 3.0 | G416 | | |
| | | | K417 | 3.1 | N92 |
| | S56 | 3.4 | D420 | | |
| | S53 | 2.8 | Y421 | | |
| | G54 | 1.9 | Y421 | | |
| | Y33 | 1.9 | L455 | | |
| | S53 | 2.1 | R457 | | |
| | S30 | 2.5 | K458 | | |
| | G54 | 2.5 | K460 | | |
| | S31 | 2.2 | Y473 | | |
| | I28 | 2.2 | A475 | | |
| | N32 | 2.6 | A475 | | |
| | Y100 | 1.6 | E484 | | |
| | G26 | 2.4 | N487 | | |
| | Y100 | 2.9 | F490 | | |
| | | | S494 | 3.3 | Y32 |

**Table S 10**: The hydrogen bonds and salt-bridges between SARS-CoV-2 RBD containing one mutation (N501Y) labeled as MT1 in the manuscript and the hNABs B38 (PDB:7BZ5).

| $\lambda = 0.22 - 0.53$ nm | | Heavy Chain | Length(Å) | SARS-CoV-2 RBD | Length(Å) | Light Chain |
|---|---|---|---|---|---|---|
| Hydrogen bond | S56 | 2.8 | G416 | | | |
| | Y52 | 3.0 | K417 | 3.1 | N94 | |
| | S56 | 1.8 | D420 | | | |
| | G54 | 1.9 | Y421 | | | |
| | | | Y453 | 3.1 | N92 | |
| | Y33 | 1.7 | L455 | | | |
| | S53 | 1.6, 2.6 | R457 | | | |
| | S30 | 2.4 | K458 | | | |
| | S31 | 2.0 | K458 | | | |
| | S31 | 1.8 | Y473 | | | |
| | I28 | 2.2 | A475 | | | |
| | N32 | 2.7 | A475 | | | |
| | G26 | 2.6 | N487 | | | |
| | R97 | 1.8, 2.0 | N487 | | | |
| | Y100 | 2.2 | F490 | | | |
| | | | Q493 | 2.5 | Y32 | |
| | | | **Y501** | 2.7 | S30 | |

**Table S 11**: The hydrogen bonds and salt-bridges between SARS-CoV-2 RBD containing one mutation (N501Y) labeled as MT1 in the manuscript and the hNAB B38 (PDB:7BZ5).

| $\lambda = 0.78 - 0.99$ nm | Heavy Chain | Length(Å) | SARS-CoV-2 RBD | Length(Å) | Light Chain |
|---|---|---|---|---|---|
| Hydrogen bond | I28 | 2.3 | A475 | | |
| | N32 | 2.6 | A475 | | |
| | G26 | 2.0 | N487 | | |
| | R97 | 1.9, 2.8 | N487 | | |
| $\lambda = 1.37 - 1.57$ nm | | | | | |
| Hydrogen bond | S53 | 2.1 | A475 | | |
| | N73 | 3.1 | S477 | | |
| | S31 | 2.9 | T478 | | |
| | S31 | 2.3 | N487 | | |
| $\lambda = 1.55 - 1.80$ nm | | | | | |
| Hydrogen bond | S31 | 2.8 | N487 | | |

**Table S 12**: The hydrogen bonds and salt-bridges between SARS-CoV-2 RBD containing three mutations (K417N/E484K/N501Y) labeled as MT3 in the manuscript and the hNAB B38 (PDB:7BZ5).

| $\lambda = 0.02 - 0.13$ nm | Heavy Chain | Length(Å) | SARS-CoV-2 RBD | Length(Å) | Light Chain |
|---|---|---|---|---|---|
| Hydrogen bond | | | R403 | 2.0, 2.1 | N92 |
| | Y58 | 2.4 | T415 | | |
| | S56 | 1.6 | D420 | | |
| | S53 | 1.8, 2.8 | Y421 | | |
| | G54 | 2.9 | Y421 | | |
| | | | Y453 | 2.6 | N92 |
| | Y33 | 2.0 | L455 | | |
| | S31 | 2.1 | Y473 | | |
| | I28 | 2.6 | A475 | | |
| | N32 | 2.0 | A475 | | |
| | G26 | 2.6 | N487 | | |
| | R97 | 1.9, 1.9 | N487 | | |
| | R97 | 3.1 | Y489 | | |
| | Y100 | 3.2 | F490 | | |
| | | | G502 | 2.0 | G28 |
| | | | Y505 | 1.9 | Q90 |
| | | | Y505 | 2.1 | L91 |
| $\lambda = 1.37 - 1.57$ nm | | | | | |
| Hydrogen bond | | | R403 | 1.9, 2.3 | N92 |
| | | | Q409 | 3.1 | Y94 |
| | Y33 | 2.1 | **N417** | | |
| | | | Y453 | 2.7 | N92 |
| | | | L455 | 1.9 | Y33 |
| | S31 | 2.0 | Y473 | | |
| | I28 | 2.4 | A475 | | |
| | N32 | 2.1 | A475 | | |
| | G26 | 2.2 | N487 | | |
| | R97 | 1.9, 3.0 | N487 | | |
| | Y100 | 2.6 | F490 | | |
| | Y100 | 2.8 | Q493 | | |
| | | | G496 | 2.3 | S30 |
| | | | Q498 | 2.7 | S67 |
| | | | T500 | 3.1 | G28 |
| | | | G502 | 2.0 | G28 |
| | | | V503 | 3.1 | Q27 |
| | | | Y505 | 2.1 | Q90 |
| | | | Y505 | 1.7 | L91 |

**Table S 13**: The hydrogen and salt-bridges between SARS-CoV-2 RBD and C102 (PDB: 7K8M) antibody.

| $\lambda = 0.02 - 0.13$ nm | Light Chain | Length(Å) | SARS-CoV-2 RBD | Length(Å) | Heavy Chain |
|---|---|---|---|---|---|
| Hydrogen bond | G92 | 2.8 | R404 | | |
| | | | G416 | 3.1 | S56 |
| | | | K417 | 1.8 | G97 |
| | | | D420 | 1.7 | S56 |
| | | | Y421 | 2.2 | G54 |
| | | | L455 | 1.8 | Y33 |
| | | | R457 | 2.2 | S53 |
| | | | K458 | 3.3 | S30 |
| | | | N460 | 2.9 | G54 |
| | | | Y473 | 1.6 | S31 |
| | | | A475 | 2.0 | I28 |
| | | | A475 | 2.1 | N32 |
| | S30 | 2.8 | N501 | | |
| | S29 | 3.1 | G502 | | |
| | Y32 | 2.0 | Y505 | | |

**Table S 14**: The hydrogen and salt-bridges between SARS-CoV-2 RBD and C102 (PDB: 7K8M) antibody.

| $\lambda = 0.10 - 0.27$ nm | Light Chain | Length(Å) | SARS-CoV-2 RBD | Length(Å) | Heavy Chain |
|---|---|---|---|---|---|
| Hydrogen bond | | | K417 | 3.6 | Y52 |
| | | | K417 | 1.7 | G97 |
| | | | D420 | 1.5 | S56 |
| | | | Y421 | 3.0 | Y33 |
| | | | Y421 | 1.8 | G54 |
| | | | Y453 | 2.4 | Y99 |
| | | | L455 | 1.9 | Y33 |
| | | | R457 | 1.8, 2.7 | S53 |
| | | | K458 | 3.0 | S31 |
| | | | K458 | 3.9 | G54 |
| | | | N460 | 3.0 | S56 |
| | | | Q474 | 3.6 | S31 |
| | | | A475 | 1.8 | I28 |
| | | | A475 | 2.5 | N32 |
| | | | Y489 | 2.7 | R94 |
| | | | Q493 | 1.6 | Y99 |

**Table S 15**: The hydrogen and salt-bridges between SARS-CoV-2 RBD and C102 (PDB: 7K8M) antibody.

| $\lambda = 0.36 - 0.60$ nm | Light Chain | Length(Å) | SARS-CoV-2 RBD | Length(Å) | Heavy Chain |
|---|---|---|---|---|---|
| Hydrogen bond | | | R457 | 2.0 | S53 |
| | | | Y473 | 2.3 | S31 |
| | | | A475 | 2.8 | I28 |
| | | | A475 | 1.9 | N32 |
| | | | S477 | 1.9 | G26 |
| | | | Y489 | 2.2 | R94 |
| $\lambda = 0.78 - 0.97$ nm | | | | | |
| Hydrogen bond | | | A475 | 2.1 | I28 |
| | | | A475 | 2.2 | N32 |
| | | | S477 | 2.2 | G26 |
| | | | N487 | 3.2, 3.4 | R94 |
| | | | Y489 | 3.1 | R94 |
| $\lambda = 1.26 - 1.52$ nm | | | | | |
| Hydrogen bond | | | N487 | 3.0, 3.1 | S31 |
| $\lambda = 1.64 - 1.87$ nm | | | | | |
| Hydrogen bond | | | S477 | 1.8, 2.8 | G54 |
| | | | N487 | 1.8, 3.3 | S53 |

**Table S 16**: The hydrogen bonds and salt-bridges between SARS-CoV-2 RBD containing one mutation (N501Y) and labeled MT1 and the hNAb C102 (PDB: 7K8M).

| $\lambda = 0 - 0.13$ nm | Light Chain | Length(Å) | SARS-CoV-2 RBD | Length(Å) | Heavy Chain |
|---|---|---|---|---|---|
| Hydrogen bond | G92 | 3.3 | R403 | | |
| | | | K417 | 2.2 | G97 |
| | | | D420 | 1.7 | S56 |
| | | | Y421 | 2.1 | G54 |
| | | | Y421 | 2.9 | G55 |
| | | | Y453 | 2.1 | Y99 |
| | | | L455 | 2.0 | Y33 |
| | | | R457 | 1.6, 2.8 | S53 |
| | | | Y473 | 1.7 | S31 |
| | | | A475 | 2.1 | I28 |
| | | | A475 | 2.1 | N32 |
| | S29 | 2.5 | G502 | | |
| | S30 | 3.4 | Y505 | | |

**Table S 17**: The hydrogen bonds and salt-bridges between SARS-CoV-2 RBD containing one mutation (N501Y) and labeled MT1 and the hNAb C102 (PDB: 7K8M).

| $\lambda = 0.05 - 0.23$ nm | Light Chain | Length(Å) | SARS-CoV-2 RBD | Length(Å) | Heavy Chain |
|---|---|---|---|---|---|
| Hydrogen bond | G92 | 2.4 | R403 | | |
| | | | K417 | 1.7 | G97 |
| | | | D420 | 1.7 | S56 |
| | | | Y421 | 2.1 | G54 |
| | | | L455 | 2.0 | Y33 |
| | | | R457 | 2.0, 2.9 | S53 |
| | | | Y473 | 1.9 | S31 |
| | | | A475 | 2.5 | I28 |
| | | | A475 | 2.3 | N32 |
| | | | Y489 | 2.7 | R94 |
| | S29 | 2.8 | G502 | | |
| | S30 | 2.2 | Y505 | | |

**Table S 18**: The hydrogen bonds and salt-bridges between SARS-CoV-2 RBD containing one mutation (N501Y) and labeled MT1 and the hNAb C102 (PDB: 7K8M).

| $\lambda = 0.15 - 0.33$ nm | Light Chain | Length(Å) | SARS-CoV-2 RBD | Length(Å) | Heavy Chain |
|---|---|---|---|---|---|
| Hydrogen bond | | | K417 | 1.8 | G97 |
| | | | D420 | 2.6 | S56 |
| | | | Y421 | 2.2 | G54 |
| | | | Y421 | 3.3 | S56 |
| | | | L455 | 1.9 | Y33 |
| | | | R457 | 2.0, 3.1 | S53 |
| | | | N460 | 2.9 | G54 |
| | | | Y473 | 2.2 | S31 |
| | | | Q474 | 3.2 | S31 |
| | | | A475 | 2.3 | I28 |
| | | | A475 | 1.9 | N32 |
| | Y32 | 3.2 | Q493 | | |
| | | | Q493 | 2.8 | Y99 |
| | G92 | 1.8 | Y505 | | |

**Table S 19**: The hydrogen bonds and salt-bridges between SARS-CoV-2 RBD containing one mutation (N501Y) and labeled MT1 and the hNAb C102 (PDB: 7K8M).

| $\lambda = 0.27 - 0.53$ nm | Light Chain | Length(Å) | SARS-CoV-2 RBD | Length(Å) | Heavy Chain |
|---|---|---|---|---|---|
| Hydrogen bond | | | K417 | 1.6 | G97 |
| | | | D420 | 1.9 | S56 |
| | | | Y421 | 2.5 | Y33 |
| | | | Y421 | 2.5 | S53 |
| | | | L455 | 1.7 | Y33 |
| | | | R457 | 1.9, 3.0 | S53 |
| | | | N460 | 2.5 | G56 |
| | | | Y473 | 1.7 | S31 |
| | | | A475 | 1.9 | I28 |
| | | | A475 | 1.9 | N32 |
| | | | A477 | 1.9,3.5 | G26 |
| | | | Y489 | 1.8, 2.9 | R94 |
| | Y32 | 2.3 | Q493 | | |
| | | | Q493 | 1.7, 3.1 | Y99 |
| $\lambda = 0.83 - 1.1$ nm | | | | | |
| Hydrogen bond | | | A475 | 2.2 | I28 |
| | | | A475 | 2.5 | N32 |
| | | | A477 | 1.9,2.8 | G26 |
| $\lambda = 0.94 - 1.18$ nm | | | | | |
| Hydrogen bond | | | Y473 | 3.2 | S31 |
| | | | A475 | 2.2 | I28 |
| | | | A475 | 2.1 | N32 |
| | | | A477 | 1.8,2.5 | G26 |

**Table S 20**: The hydrogen bonds and salt-bridges between SARS-CoV-2 RBD containing three mutations (K417N/E484K/N501Y)and labeled MT3 and the hNAb C102 (PDB: 7K8M).

| $\lambda = 0.02 - 0.13$ nm | Light Chain | Length(Å) | SARS-CoV-2 RBD | Length(Å) | Heavy Chain |
|---|---|---|---|---|---|
| Hydrogen bond | | | T415 | 3.0 | S56 |
| | | | **N417** | 2.0, 3.6 | Y33 |
| | | | **N417** | 2.8 | G97 |
| | | | D420 | 1.8 | S56 |
| | | | Y421 | 2.5 | G54 |
| | | | Y421 | 2.8 | G55 |
| | | | Y421 | 3.1 | S56 |
| | | | R457 | 1.6, 2.9 | S53 |
| | | | R458 | 2.6 | S31 |
| | | | N460 | 1.9 | S56 |
| | | | Y473 | 1.8 | S31 |
| | | | Q474 | 3.3 | S31 |
| | | | A475 | 2.3 | I28 |
| | | | A475 | 1.8 | N32 |
| | | | Y489 | 2.9 | R94 |
| | S29 | 3.1 | G502 | | |
| | Y32 | 3.4 | Y505 | | |
| $\lambda = 0.7 - 0.9$ nm | | | | | |
| Hydrogen bond | | | **N417** | 2.7 | Y33 |
| | | | D420 | 1.8 | S56 |
| | | | Y421 | 1.9 | G54 |
| | | | Y453 | 2.7 | Y99 |
| | | | L455 | 1.6 | Y33 |
| | | | R457 | 2.2 | S53 |
| | | | Y473 | 1.8 | S31 |
| | | | A475 | 2.1 | I28 |
| | | | A475 | 1.9 | N32 |

### 4 Class II antibodies

Below we present the amino acid residues that are interacting between the RBD (in its WT and mutated versions) and with Class II antibodies P2B-2F6 and C144.

**Table S 21**: The hydrogen bonds and salt-bridges between SARS-CoV-2 RBD and P2B-2F6 (PDB: 7BWJ) antibody

| $\lambda = 0 - 0.14$ nm | Heavy chain | Length(Å) | SARS-CoV-2 RBD | Length(Å) | Light chain |
|---|---|---|---|---|---|
| Hydrogen bond | Y27 | 1.9 | G447 | | |
| | S31 | 1.9 | Y449 | | |
| | S30 | 1.8 | N450 | | |
| | H54 | 1.9 | N450 | | |
| | | | G482 | 3.3 | Y34 |
| | R112 | 1.8, 3.2 | E484 | 2.4 | N33 |
| | | | E484 | 2.6 | Y34 |
| Salt bridge | R112 | 2.0, 3.7 | E484 | | |
| $\lambda = 0.62 - 0.87$ nm | | | | | |
| Hydrogen bond | R112 | 1.7, 2.4 | E484 | | |
| | | | G482 | 1.9 | Y34 |
| Salt bridge | R112 | 0.8, 1.7 | E484 | | |
| $\lambda = 0.98 - 0.76$ nm | | | | | |
| Hydrogen bond | R112 | 2.6, 3.1 | E484 | | |
| | | | G482 | 3.4 | Y34 |
| Salt bridge | R112 | 3.1 | E484 | | |
| $\lambda = 1.15 - 1.4$ nm | | | | | |
| Hydrogen bond | | | E484 | 2.3 | Y34 |

**Table S 22**: The hydrogen bonds and salt-bridges between SARS-CoV-2 RBD and C144 (PDB: 7K90) antibody.

| $\lambda = 0 - 0.12$ nm | Light Chain | Length(Å) | SARS-CoV-2 RBD | Length(Å) | Heavy Chain |
|---|---|---|---|---|---|
| Hydrogen bond | | | K417 | 3.0 | Y100 |
| | | | E484 | 1.7, 2.2 | S53 |
| | | | E484 | 1.9 | G54 |
| | | | E484 | 2.1, 2.3 | S56 |
| | | | E484 | 3.1 | R100 |
| | | | Y489 | 1.7 | D100 |
| | | | F490 | 1.8, 3.2 | R100 |
| | | | L492 | 2.6 | R100 |
| | | | Q493 | 1.8, 2.6 | R100 |
| | | | S494 | 3.5 | S30 |
| | | | S494 | 1.9, 2.2, 2.8, 3.2 | N31 |
| $\lambda = 0.32 - 0.48$ nm | | | | | |
| Hydrogen bond | | | E484 | 1.8 | S53 |
| | | | E484 | 3.5 | S56 |
| | | | E484 | 1.9, 2.7 | R100 |
| | | | Y489 | 1.9, 2.5 | D100 |
| Salt bridge | | | E484 | 3.5 | R100 |
| $\lambda = 0.53 - 0.75$ nm | | | | | |
| | | | F486 | 3.2 | S56 |
| $\lambda = 0.93 - 1.14$ nm | | | | | |
| Hydrogen bond | | | F486 | 1.8 | G54 |

**Table S 23**: The hydrogen bonds and salt-bridges between SARS-CoV-2 RBD containing three mutations (K417N/E484K/N501Y) and labeled MT3 and the hNAbs C144 (PDB: 7K90).

| $\lambda = 0 - 0.12$ nm | Light Chain | Length(Å) | SARS-CoV-2 RBD | Length(Å) | Heavy Chain |
|---|---|---|---|---|---|
| Hydrogen bond | | | Y449 | 2.8 | G26 |
| | | | **K484** | 2.6 | S55 |
| | Y91 | 3.0 | F486 | | |
| | | | N487 | 3.0 | D100 |
| | | | F490 | 2.0, 2.8 | R100 |
| | | | L492 | 2.6 | R100 |
| | | | Q493 | 1.9, 2.4 | R100 |
| | | | S494 | 2.1, 2.1 | N31 |
| | | | Q498 | 2.9 | G26 |
| $\lambda = 0.7 - 0.9$ nm | | | | | |
| Hydrogen bond | | | **N417** | 2.3 | Y100 |
| | | | Y489 | 2.4 | R100 |
| | | | Q493 | 2.6 | R100 |

### 5 Class III antobodies

Below we present the amino acid residues that are interacting between the RBD and with Class III antibody C135.

**Table S 24**: The hydrogen bonds and salt-bridges between SARS-CoV-2 RBD and C135 (PDB: 7K8Z) antibody.

| $\lambda = 0.07 - 0.29$ nm | Heavy Chain | Length(Å) | SARS-CoV-2 RBD | Length(Å) | Light Chain |
|---|---|---|---|---|---|
| Hydrogen bond | Y98 | 3.4 | T345 | | |
| | | | T345 | 2.4 | Y94 |
| | R55 | 2.1 | N437 | | |
| | D53 | 1.9 | N440 | | |
| | N56 | 2.6 | N440 | | |
| | S96 | 1.9 | V445 | | |
| | | | Y451 | 3.2 | W-32 |
| $\lambda = 0.26 - 0.55$ nm | | | | | |
| Hydrogen bond | | | T345 | 1.9 | N92 |
| | R55 | 1.9, 2.8 | S438 | | |
| | D53 | 1.8 | N440 | | |
| | N56 | 2.2 | N440 | | |
| | S96 | 2.9 | V445 | | |
| $\lambda = 0.34 - 0.61$ nm | | | | | |
| Hydrogen bond | | | T345 | 1.9 | N92 |
| | R55 | 2.2, 2.3 | S438 | | |
| | D53 | 1.7, 2.6 | N440 | | |
| | N56 | 2.2 | N440 | | |
| | S96 | 1.9 | V445 | | |
| $\lambda = 0.36 - 0.68$ nm | | | | | |
| Hydrogen bond | | | T345 | 1.9 | N92 |
| $\lambda = 0.59 - 0.86$ nm | | | | | |
| Hydrogen bond | | | T345 | 2.0 | N92 |
| | S96 | 1.9 | V445 | | |
| $\lambda = 0.91 - 1.23$ nm | | | | | |
| Hydrogen bond | | | T345 | 1.7, 2.2, 3.1 | E50 |
| | S96 | 2.3,3.2 | V445 | | |
| $\lambda = 1.05 - 1.35$ nm | | | | | |
| Hydrogen bond | G97 | 1.8 | T345 | | |
| | | | R346 | 1.7, 2.2 | E50 |
